# The *in vitro* dynamics of pseudo-vascular network formation

**DOI:** 10.1101/2023.11.02.565264

**Authors:** Mariam-Eleni Oraiopoulou, Dominique-Laurent Couturier, Ellie V. Bunce, Ian Cannell, Monika Golinska, Gregory Hannon, Vangelis Sakkalis, Sarah E. Bohndiek

## Abstract

Pseudo-vascular network formation capacity *in vitro* is considered a key characteristic of vasculogenic mimicry. While many cancer cell lines are known to form pseudo-vascular networks, little is known about the spatiotemporal dynamics of these formations. Here, we present a framework for monitoring and characterising the dynamic formation and dissolution of pseudo-vascular networks *in vitro*. The framework combines time-resolved optical microscopy with open-source image analysis for network feature extraction and statistical modelling. The framework is demonstrated by comparing diverse cancer cell lines associated with vasculogenic mimicry, then in detecting response to drug compounds proposed to affect formation of vasculogenic mimics. Dynamic datasets collected were analysed morphometrically and a descriptive statistical analysis model was developed in order to measure stability and dissimilarity characteristics of the pseudo-vascular networks formed. Melanoma cells formed the most stable pseudo-vascular networks and were selected to evaluate the response of their pseudo-vascular networks to treatment with axitinib, brucine and tivantinib. Our framework is shown to enable quantitative analysis of both the capacity for network formation, linked vasculogenic mimicry, as well as dynamic responses to treatment.

## Introduction

Angiogenesis, the growth of new blood vessels, has long been recognised as an important contributor to solid tumour progression, yet therapeutic targeting has often shown limited efficacy and short-term benefit for the patient [1]. Emerging from this surprising outcome is the knowledge that cancer vascularisation proceeds through mechanisms other than angiogenesis, which are genotypically and phenotypically distinct. Among them, yet outside of the endothelial context, vasculogenic mimicry (VM) is characteristic of aggressive cancers, first observed over two decades ago in human melanoma [2], with an epidemiologic incidence ranging from 5.25% in melanoma to 65.9% in glioblastoma [3]. Furthermore, radiotherapy and chemotherapy can lead to VM-related epigenetic transition of cancer cells in treated tumours [4] and VM-rich tumours are associated with therapeutic resistance [5, 6].

Unlike angiogenic mechanisms, where new blood vessels are formed from pre-existing vessels to compensate oxygen and nutrient deficiency in the growing tumour mass, VM vessels have the unique characteristic that they arise solely from cancer cells that trans-differentiate into an endothelial-like phenotype [5, 6]. Specifically targeting VM-rich tumours may therefore benefit from a dual anti-vascular and chemotherapeutic approach.

The core characteristic typically used to define VM *ex vivo* is the presence of vessels in Periodic acid–Schiff (PAS) stained sections that lack counter-staining for endothelial markers (e.g. CD31/CD34) [5, 7, 8], and ideally contain red blood cells to demonstrate functionality. Cancer cell lines where VM has been identified *ex vivo* have been shown to organise into interconnected polygonal cellular networks *in vitro* akin to those formed by endothelial cells [7, 9]. Hence the formation of pseudo-vascular networks has further been defined as a core characteristic of VM competence. A range of other VM markers have also been identified, however, these do not exclusively indicate VM; for example, poorer overall tumour oxygenation *in vivo* [10, 11] or the expression of VE-cadherin (or CD144) and matrix metalloproteinase-2 (MMP-2), among other extracellular matrix factors [12].

Although typical core characteristics of VM *ex vivo* and *in vitro* are becoming more generally accepted, little is known about VM as a pathophysiological vascularisation mechanism or its evolution over time. Most studies are undertaken at a single time point, using protocols that often vary between studies making comparisons between cancer types or cell lines challenging [13]. Imaging data from *ex vivo* sections or *in vitro* cultures are rarely subjected to quantitative object extraction, meaning readers are left to infer network characteristics from optical micrographs. Furthermore, the pseudo-vascular networks formed are often compared visually to those formed by non-cancerous endothelial cell lines, which is inappropriate as mechanistically these exhibit angiogenic phenotypes [14–16]. The high degree of variation between studies, along with inconsistency among different reports, has led to controversy in the field of VM, particularly for *in vitro* studies [17]. In addition, different morphologies have been observed *in vitro* [9] that are not well distinguished *in vivo*, heightening the debate. Taken together, improved implementation and quantification of *in vitro* VM assays is now important to advance the field and increase reproducibility of research.

To more thoroughly describe the *in vitro* VM phenotype, we applied an experimental framework based on an endothelial tube assay, where cultures are monitored dynamically as they form and dissolve pseudo-vascular networks. Combining optical microscopy data with open-source Fiji software, we analysed morphometrically the spatiotemporal dynamics of the networks formed by a range of different cell lines representing brain, breast, head and neck, and skin cancer types, that have been previously reported to demonstrate VM [5]. A descriptive statistical model was implemented to compare the morphology and stability of the pseudo-vascular networks before applying a range of concentrations of therapeutics that have been reported to have VM-related mechanisms of action. We identify relevant features of the networks that can be used to compare different cell lines and drug responses, demonstrating the potential of our methodology to evaluate VM-competence *in vitro*.

## Methods

### Cell Cultures

U87MG (HTB-14™, ATCC, UK) cells were used to represent human high-grade brain cancer. MDA-MB-231 (HTB-26™, ATCC, UK) and 4T1 (CRL-2539™, ATCC, UK) cell lines were used as mammary breast carcinomas of human and murine origin respectively. A VM-competent 4T1 derivative clone (4T1-T) reported in previous work [7] was also tested. Murine MOC1 (CVCL_ZD32, Kerafast, USA) and MOC2 (CVCL_ZD33, Kerafast, USA) cell lines were used to represent head and neck squamous carcinoma. B16-F10 (CRL-6475™, ATCC, UK) murine cell line was used to represent melanoma skin cancer.

All cells were cultured in standard cell culture conditions in the absence of any antibiotics with 10% fetal bovine serum (FBS, Gibco, USA) supplementation. The media used were Roswell Park Memorial Institute 1640 medium (RPMI, Gibco, USA) for the 4T1 and 4T1-T cell lines or Dulbecco’s Modified Eagle medium (DMEM, Gibco, USA) for the MDA-MB-231, U87MG and B16-F10 cell lines. MOC1 and MOC2 cell lines were grown in DMEM supplemented with 10% M-FBS (Gibco, Mexico). All cell lines were profiled using short tandem repeats (STR) genotyping to validate their status compared to the original archived profiles where available and were >95% authenticated. For all cell lines, cell passages used in the experiments were limited to 3 above the starting cell passage.

### Doubling Time Assay

Cells were cultured in 48-well plates in a range of initial cell seeding densities (from 500 to 10^4^ cells in 500μl of medium per well) to assess their doubling times and growth curves. Cell growth was monitored for up to 72hrs using automated Incucyte S3 live cell analysis system (Sartorius, Germany). Phase contrast images were captured every 3hrs in 4x magnification throughout the incubation time. Growth rate estimations were extracted based on confluence masks of the phase contrast images and were analysed for each cell seeding density. In each experiment, each condition was performed in triplicate and all experiments were repeated at least two times.

### 2D Drug Screening Assay

In a 96-well black clear-bottom plate (Corning, USA), 2500 cells in 100μl of medium were seeded per well and incubated overnight to allow adhesion. The cells were treated with a range of 0.5-120μM for axitinib (stock solution 50mM, Generon, France), 6-120μM for brucine (stock solution 10mM, Generon, France) and 1.5-120μM for tivantinib (stock solution 15mM, ARQ-197, Generon, France). As all drug agents were diluted in dimethyl sulfoxide (DMSO, Sigma Aldrich, USA), treatment with 0.1% DMSO was used as a control. The plate was then incubated for up to 72hrs post-treatment in the Incucyte S3 (Sartorius, Germany) under the same acquisition parameters as above.

At the end of the time course, each well was treated with 1:1 CellTiter-Glo (Promega, UK) and incubated for 10mins at room temperature. Luminous signal was detected with CLARIOstar® (BMG LABTECH, Germany) and the cell viability was estimated to confirm inhibition effect of the agents used and validate the Incucyte data.

### Tubular Network Formation Assay

A summary of the experimental pipeline used for the purposes of this study is shown in Figure 1**Error! Reference source not found.**A. A layer of basement membrane extract (BME, Cultrex, UK), of Type R1 (reduced growth factor, RGF BME) and protein content greater than 9, was first coated in a 24-well plate and allowed to solidify for 1hr. Cells were trypsinized and resuspended in endothelial cell growth basal medium-2 (EBM-2, Lonza, Switzerland). Cell seeding density into wells was 0.6 × 10^5^ cells in 300μl medium, which was experimentally confirmed as the optimal initial cell concentration for all the cell lines tested based on the assays above.

**Figure 1:**
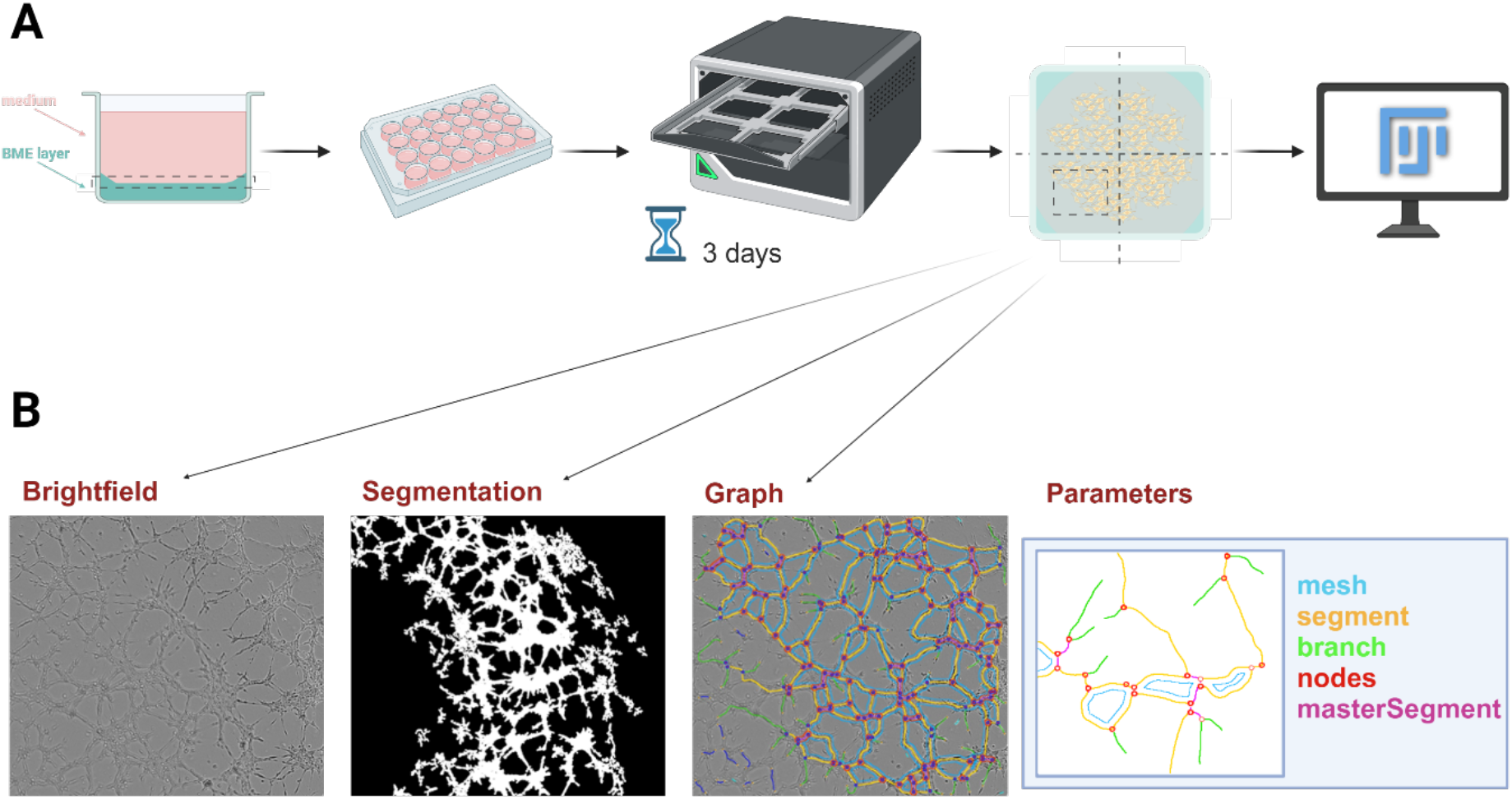
Schematic overview of the framework applied to evaluate pseudo-vascular network dynamics in cancer cell lines. A. Experimental steps (from left to right): cell/ medium component and basement membrane extract scaffold on each well of a 24-well plate, incubation and monitoring by phase contrast microscopy, and subsequent image analysis using Fiji. B. Image analysis pipeline (from left to right): the brightfield image is processed into a binary network mask and Angiogenesis Analyzer applied to extract a graph segmentation of a representative pseudo-vascular network as shown, along with vectorial object features.

Where applicable, cells were treated after seeding with axitinib (6.25, 12.5, 25, 50, 125μM, Generon, France), brucine (12.5, 25, 50, 100μM, Generon, France) or tivantinib (3.75, 7.5, 15, 25μM, Generon, France). DMSO 0.01% was used for the control untreated wells.

Each plate was monitored for up to 96hrs in the Incucyte S3 and phase contrast photographs were captured every 1hr in 4x magnification across the 4 quadrants of the well. In each experiment, each condition was performed in triplicate and all experiments were repeated at least three times. Exemplar videos are provided as supplementary materials (Supplementary Videos 1 – 7).

### Image Analysis

The image analysis pipeline is summarized in Figure 1B. Incucyte software (v.2022A, Sartorius, Germany) was used to export all phase contrast images and videos. A manual quality check was applied on the acquired images to disregard medium condensation and/ or out of focus images. For each well, measurements in the four quadrants were defined as technical replicates. For each of the cell lines under study, a time window was defined for analysis based on the overall pseudo-vascular network evolution, including prior timepoints examined in the literature for the same cell line.

All technical replicates were segmented using the open-source Fiji image analysis software [18]. To avoid artifacts from the meniscus of the BME, each of the technical replicate images was cropped for a central area of 1000×1000 pixels. Each pixel was set at 2.82 μm × 2.82μm based on the Incucyte acquisition parameters. Contrast enhancement of ‘Enhance Contrast’ function (0.35 saturated) was applied to clearly segment the tube formations. Images were then transformed to RGB 8-bit format and inputted into the Fiji plugin Angiogenesis Analyzer [19]. The pseudo-vascular network images were skeletonized by the plugin to be transformed into mesh networks, which allowed their parameterisation using a total of 20 vectorial objects (network features, Supplementary Table 1) defined by the plugin.

### Statistical Analysis

For the growth curves of each cell line, linear regression analysis was performed using GraphPad Prism 9.3.1 (GraphPad, USA) using the quantification of phase contrast images from the Incucyte software (v.2022A, Sartorius, Germany) accounting for cell confluence. Spearman’s rank correlation was used to measure the level of dependence between physiological variables of the different cell lines, such as doubling time to tube formation time window, and the time window of pseudo-vascular network formation. To analyse the tubular network formations, several analyses were performed.

First, for each cell line, the average of the four technical replicates within the well was calculated for each of the 20 vectorial objects identified by the Angiogenesis Analyzer software. To examine the dynamics of the tubular networks in each well (*i*), the relationship between the object average (*y_i_*) and time (*t_i_*) was then modelled by the following third-degree polynomial model:

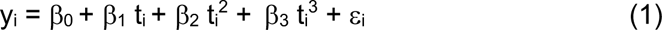

where the *β* are fit parameters and *ε_i_* denotes the error term. Three estimators were considered: linear regression (function *lm* of the *stats* R package), robust mixed-effect linear regression with 90% efficiency (function *lmeRob* of the *robust* R package) and quantile regression (function *rq* of the *quantreg* R package). The first and second estimators model the mean object value given time, while the third one models the median object value given time. The first is sensitive to outliers while the second and third are more robust to them. As results were found to be consistent between the models, the output from linear regression is displayed unless otherwise stated.

Next, to compare the results obtained for different cell lines, dissimilarity scores were used. As the time range of pseudo-vascular network formation and disappearance varies between different cell lines (e.g. 8 to 12hrs for MOC1, and 17 to 29hrs for B16-F10), the cell line time windows were first standardized using the following transformation:

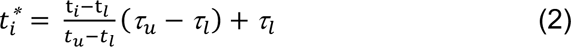

where: t*_i_* and 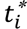, denote the original and standardized time corresponding to the *i*^th^ well of the cell line of interest; *t_u_* and *t_l_* denote the upper- and lower-time boundaries of the cell line of interest; *τ_u_* and *τ_l_* denote the upper and lower overall time range of interest.

As the vectorial objects also have very different values, we then normalized the object data using z-scores for each object:

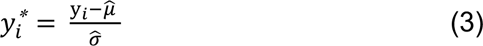

where *y_i_* and 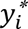, denote the original and normalized object value corresponding to the *i*^th^ well, while 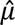 and 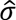. denote the average and standard deviation of all observations of the object of interest for that cell line and timepoint, so that transformed data have mean 0 and unit variance for each morphological object.

To obtain dissimilarity scores per vectorial object, we then defined the integrated discrepancy between all pairs of cell lines on the time- and object-transformed scales [obtained by Equation (2) and (3)]. For cell line pair (*j,k*), the integrated discrepancy *d_j,k_* for a given object is then given by:

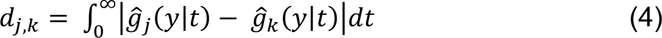

where 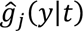 and 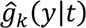, denote the estimated relationship between the object and time described in Equation (1) for cell lines *j* and *k*. The dissimilarity scores of the *f*^th^ object was then obtained by summing the integrated discrepancy of all cell line pairs. Inference for the dissimilarity scores was obtained by defining 95% percentile intervals of 4000 bootstrap dissimilarity scores obtained by means of a hierarchical bootstrap in which both wells and technical replicates within wells were sampled with replacement, and scores were defined by re-estimating Equation (4) based on fits of (1) on the transformed data described in Equations (2) and (3) for each bootstrap sample.

Finally, to compare the stability of the tubular networks between cell lines, a spread score was defined as the maximum expected value minus the minimum expected value at a given time, i.e.:

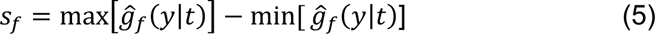

where 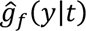 denotes the estimated relationship between object (*f*) and time (*t*) described in Equation (1) using the standardized object data from Equation (3). The stability score of each cell line was then obtained by taking the sum of the spread scores of all objects of the cell line of interest. Inference for the stability scores was obtained by defining 95% percentile intervals of 4000 bootstrap stability scores obtained by means of a hierarchical bootstrap defined similarly as the one described in Equation (5). Sensitivity analyses considering alternative stability scores (like integrated discrepancy with a horizontal line positioned at the mean object level of the cell line of interest) led to the same conclusions.

## Results

### Pseudo-vascular network growth is transient and correlates to cell growth

The doubling time measurements show that the growth rates of the two human cell lines, U87MG and MDA-MB-231 are quite similar, while all the murine cell lines appear to be more proliferative (Table 1). Pseudo-vascular tubular network formation has generally been considered at only a single timepoint in prior studies [7, 14, 16, 20]. Contrary to the pseudo-vascular networks arising from endothelial cells that are stable, characterised by a meshed 3D architecture with stable features [16], the cancer cells interrogated here are highly dynamic.

**Table 1.**
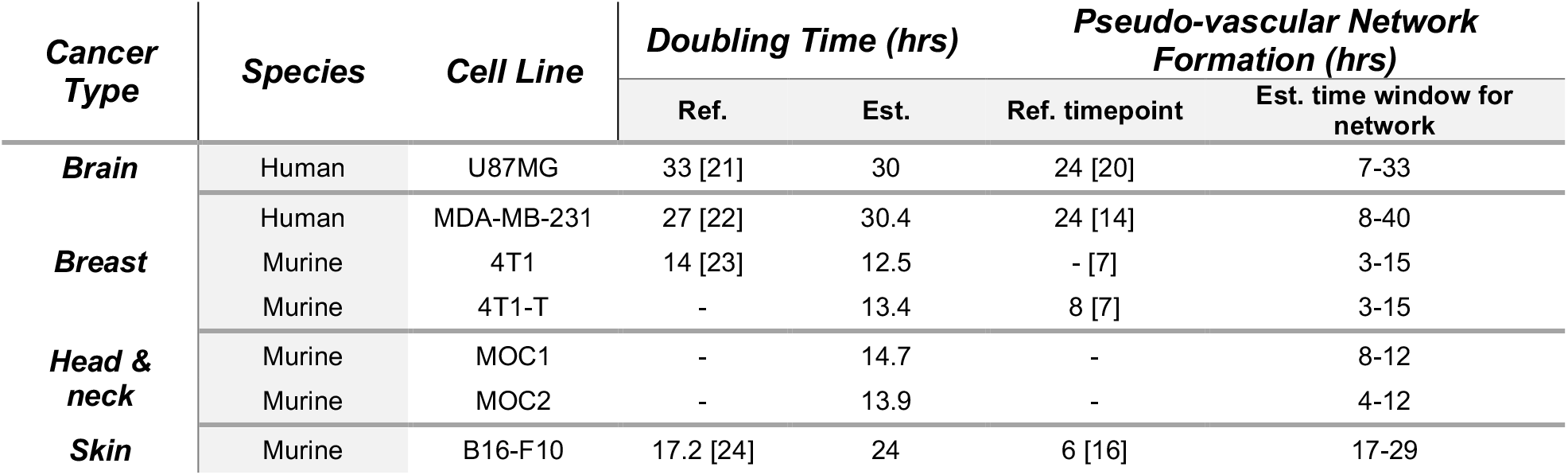
Growth metrics for each cell line compared to the literature. Error bars given show the standard error on the mean across experiments.

Our findings indicate that pseudo-vascular network formation is transient, forming prior to and disappearing after the singular timepoint typically reported, however, the time window during which such a network can be observed is different for each cancer cell line (Figure 2). As expected, the more proliferative cell lines show a shorter time window for network formation, but also a more transient network formation (Table 1). The pseudo-vascular networks formed by the slower growing U87MG and MDA-MB-231 cell lines qualitatively demonstrate a more consistent morphology across the time course. The cell line doubling time and the time window of pseudo-vascular network formation show a strong dependence level (Spearman’s rho = 0.84).

**Figure 1.**
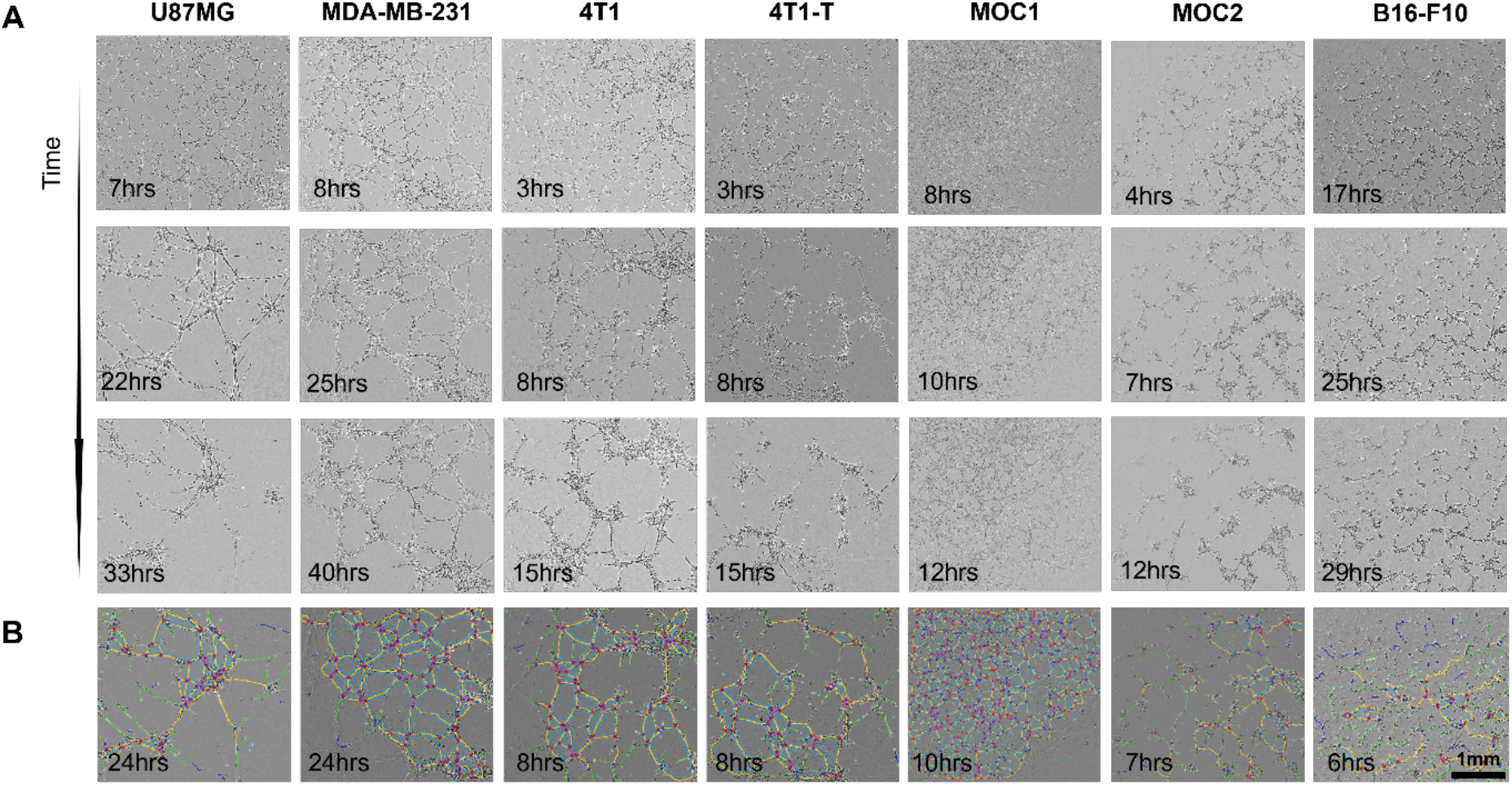
Qualitative depiction of pseudo-vascular network dynamics and single time-point graph outputs. Each column refers to a different cell line. A. Optical micrographs for an early, an intermediate and a later timepoint are depicted for each cell line, based on the respective time window over which the networks are formed. B. The Fiji graph outputs are shown at the reference timepoint (see also Table 1). Notice that for the B16-F10, a mature pseudo-vascular network has yet to be formed at this point. Scale bar is set to 1mm.

### Quantitative morphological analysis of the pseudo-vascular networks shows spatiotemporal diversity between different cancer cell lines

The BME scaffold provided in each well triggers the cells to undergo processes that are related to elongation, migration, adhesion, aggregation and division, forming pseudo-vascular networks [25] provided they have this intrinsic capacity. To gain a quantitative understanding of the structural variety observed in the evolution of the pseudo-vascular networks formed by the different cell lines, we applied the simple and precise Angiogenesis Analyzer tool designed to characterize meshed and/or branched structures of endothelial networks [19].

Before applying such quantitative analysis to these data, it is necessary to account for the effect of the meniscus (Figure 3A). When seeded in BME-coated wells, the cells tend not to colonize the boundaries of the curved meniscus (Figure 3B); the phenomenon has previously been characterized as a result of the heterogeneous viscoelastic substrate properties at the edges of the well that prohibit the cell elongation, interrupting tube formation [19]; lower cellularity is also observed at the periphery. We therefore segmented out regions from our images to exclude the meniscus prior to analysis. It is also necessary to consider potential artefacts that could bias the readout from some vectorial objects. For example, isolated structures could potentially be formed from cells that remained unattached to the pseudo-vascular network or even non-cellular components, such as macromolecules of the ECM, giving artefactual results from automated analysis. Two objects are sensitive to these isolated structures, “number of isolated segments” and “number of pieces” (Supplementary Table 1) and were found to display high variance (Figure 3C) with output outliers. Given the observed variance in these data compared to other objects, they were excluded from further analysis.

**Figure 3.**
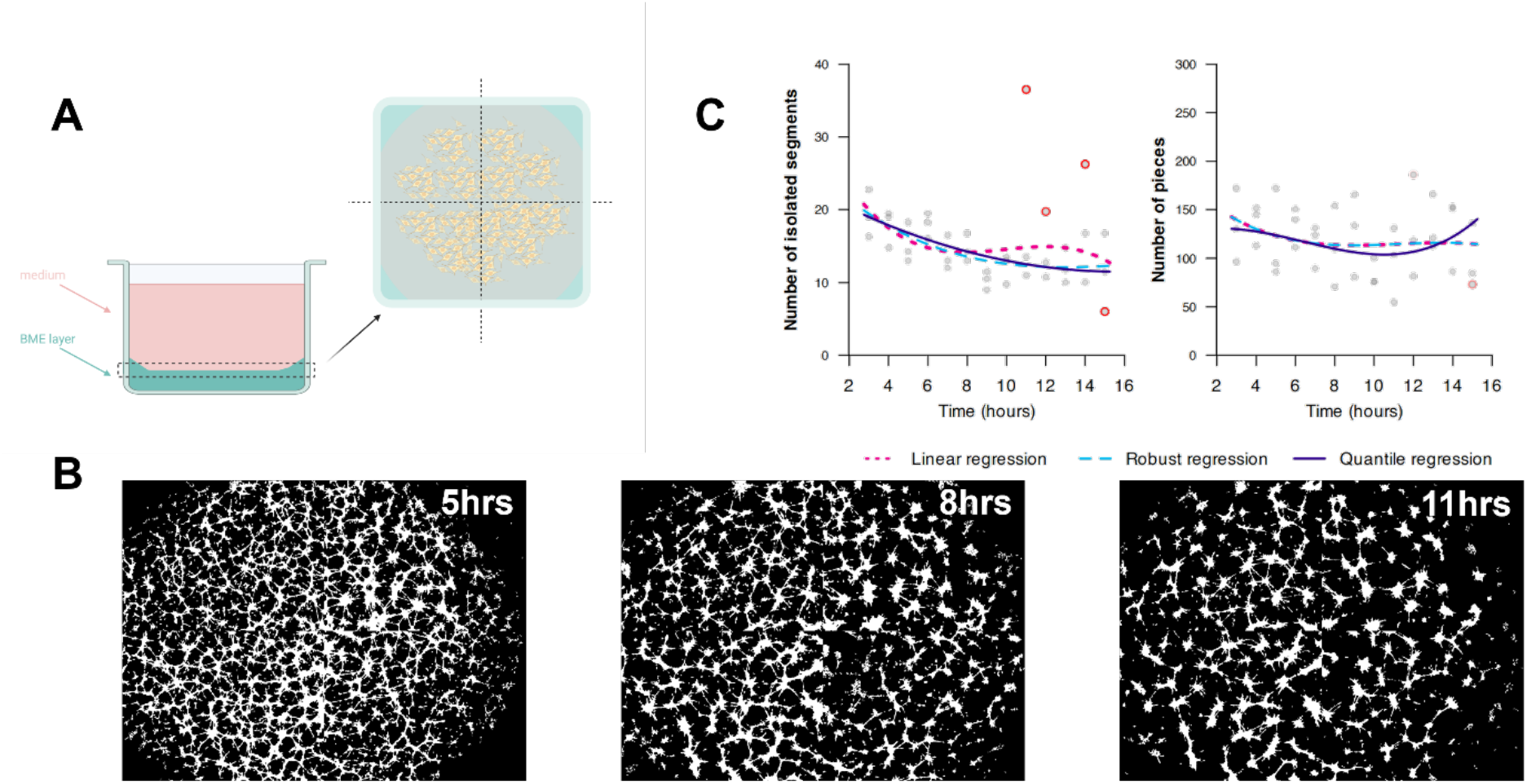
The effect of the meniscus on imaging data. A. Illustration of the meniscus effect within a cell culture well. B. Binary masks of whole well with 4T1-T cells forming tubular structures for 3 representative timepoints. Notice that the cellularity of the periphery of the well is low for all timepoints shown. C. For two morphological objects (number of isolated segments and pieces) for the 4T1-T cell line, average object value (y-axis) versus time (x-axis) is shown for each well (grey dots). Coloured lines correspond to the fit of the third-order polynomial linear model estimated by means of three estimators: the linear regression (dotted lines), the robust regression (dashed lines) and he quantile regression (solid line). Coloured points correspond to wells with outliers. based on the object-to-time relationship fitted by means of linear regression. Inference was obtained by means of a hierarchical bootstrap. B. Vectorial objects are shown in a graph. C. Spearman correlation matrix of the 4000 bootstrap dissimilarity scores aiming to measure the level of association between objects by considering similarities in dissimilarity scores.

We initially examined the full set of 20 vectorial objects output from Angiogenesis Analyzer as an average across the time course. While some cell lines show comparatively similar values for many objects over the time course (e.g. U87MG and MDA-MB-231, Supplementary Figure 1a, b), others highly dynamic and distinct (Supplementary Figure 1c-g). We can start to learn about the overall structure and stability of the pseudo-vascular networks using these parameters (Figure 4B). For example, the higher the number of extremities within the analysed area, the more disconnected the cells are, resulting in fewer tubular structures. The number of extremities is very high at the start and end of the time course for all cell lines, which helps to define the estimated time window for the persistence of the network. The number of the branches is rather stable for the U87MG, MDA-MB-231, 4T1, 4T1-T and B16-F10 cell lines, but not for MOC1 and MOC2, indicating a more dynamic network for the latter two.

**Figure 4.**
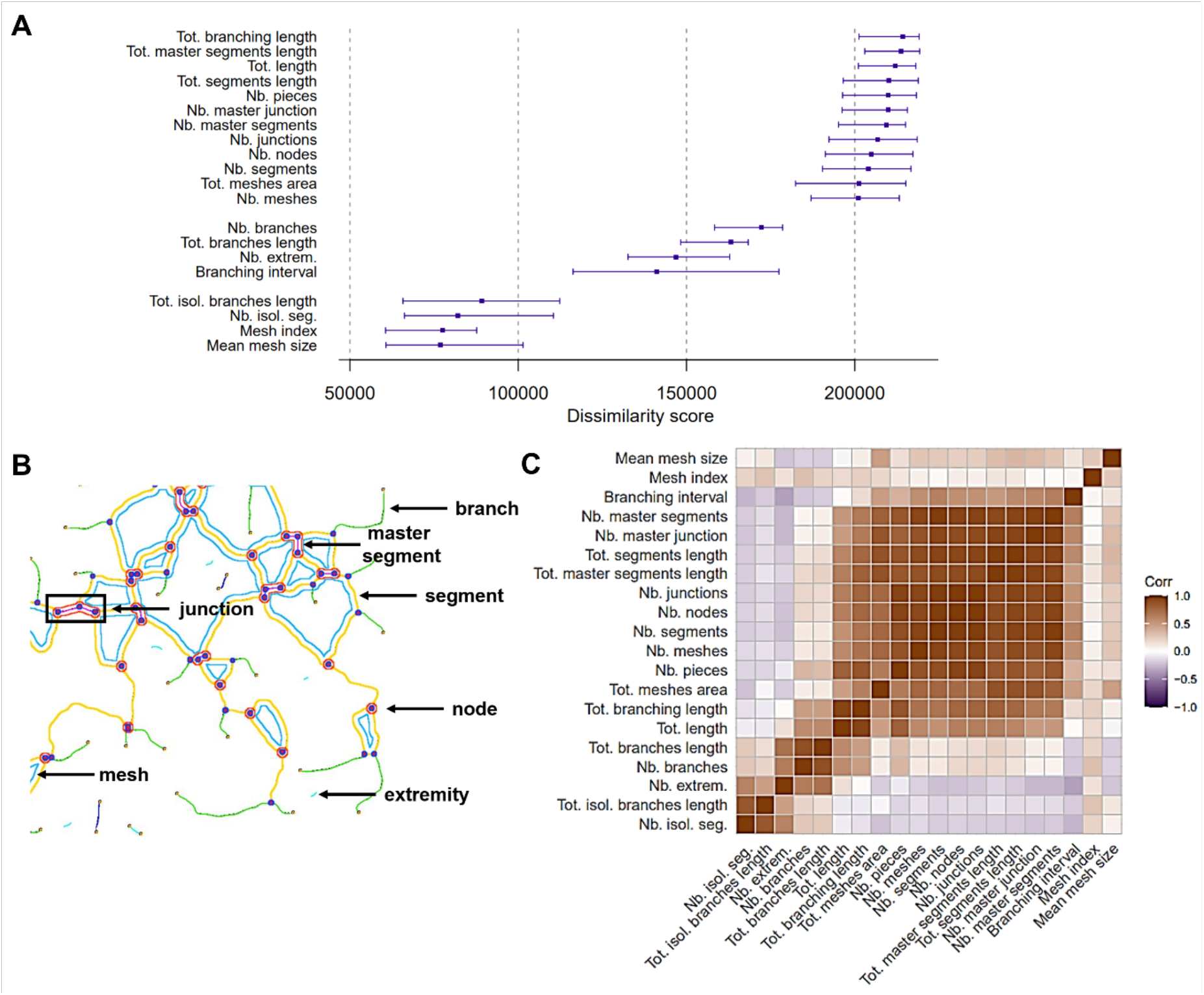
Dissimilarity scores across all cell lines for the vectorial objects extracted. A. Point and interval estimates of dissimilarity scores (x-axis) per object (y-axis) defined Across all cell lines examined, the B16-F10 melanoma cell line appears most stable. U87MG glioblastoma cells are similar but show a larger spread in some parameters, namely the number of isolated segments and total isolated branches length. While the qualitative examination of MDA-MB-231 appears relatively stable, quantitative analysis shows that their network is actually relatively unstable. The poor stability performance of the MDA-MB-231 pseudo-vascular networks primarily derives from the mesh index and mean mesh size (Supplementary Figures S1 and S2). Notably, although the aforementioned vectorial objects were not found to be good discriminators between cell lines (Figure 4A), MDA-MB-231, along with the other unstable cell line MOC2, have remarkably large mean mesh size compared to the other cell lines, which may indicate a different underlying mechanism of network development.

Interestingly, the average size of the meshes formed from all networks is similar independent of the timepoint; this can also be observed in the respective values of mesh index and mean mesh size, where only subtle differences can be observed between cell lines. Unlike the size of the meshes formed, the total mesh area, in line with the number of the meshes, confirms that MDA-MB-231, B16-F10, 4T1-T and U87MG have most cells organized into mesh components for most of the time that the networks develop, unlike the MOC1, MOC2 and 4T1 cell lines.

### Vectorial objects enable a quantitative comparison of pseudo-vascular network dynamics across different cell lines

The capability of cell lines to engage in VM is typically assessed by their ability to form a pseudo-vascular network *in vitro* [9], however, the time windows and the range of outcome values for each vectorial object of the different cell lines studied vary widely. To examine to what extent the different morphological descriptors can discriminate the pseudo-vascular networks formed by the different cell lines, we defined dissimilarity scores for each vectorial object based on the discrepancy between the dynamic fits of all pairs of cell lines (Figure 4A). The objects related to the mean size of the meshes and mesh index showed the least dissimilarity (i.e. highest similarity) between cell line fits (low value score). Conversely, the number of the meshes and the total mesh area, as well as the various length objects, were more effective at discriminating the networks.

Correlations between the objects according to their dissimilarity scores (Figure 4C) measures association between objects according to their relationship with cell lines and indicates the presence of 3 to 4 groups of highly correlated objects. Interestingly, the clustering of vectorial objects does not differ between the dissimilarity and correlation analyses, indicating that vectorial objects that show high dissimilarity scores and are correlated can serve as good morphological discriminators for pseudo-vascular networks. Based on these principles, we can narrow down the vectorial objects that are considered, avoiding redundancy. Based on these findings, we selected a subset of vectorial objects in each dissimilarity score group, 3 high (total network length, total mesh area and number of meshes), 2 average (number of branches and number of extremities) and 1 low (mesh index) discriminators, to demonstrate pseudo-vascular network dynamics (Figure 5). Our modelling approach enabled the descriptive comparison of vectorial objects of the pseudo-vascular networks formed by the different cell lines over time.

**Figure 5.**
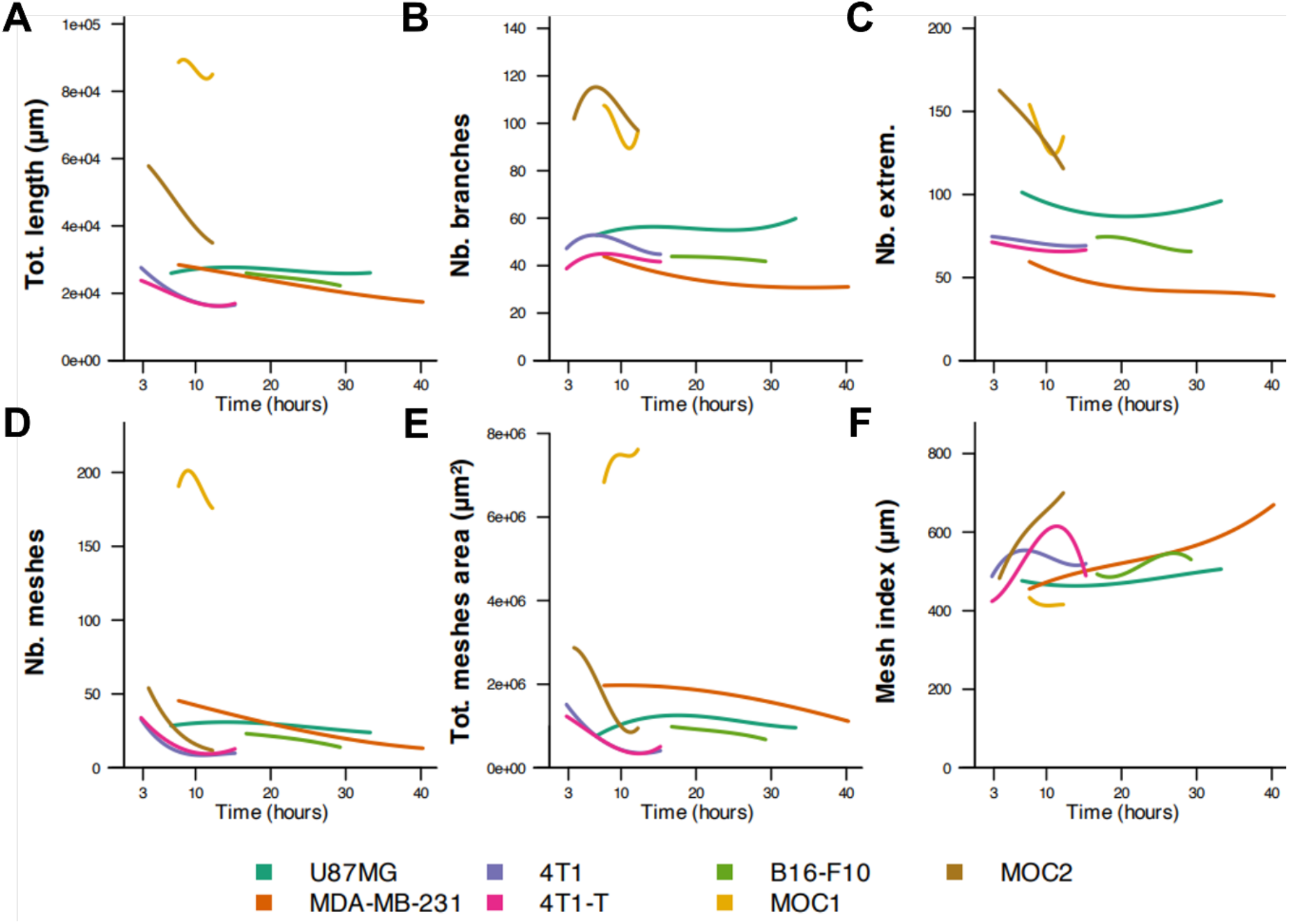
Quantitative analysis of pseudo-vascular network dynamics comparing all cell lines. Fit of the relationship between vectorial objects (y-axis) and time (x-axis) according to linear regression estimator per cell line (coloured lines) for 6 morphological objects that can serve as discriminators between the networks formed by the different cell lines. Namely, total network length (A), number of branches (B), number of extremities (C), number of meshes (D), total mesh area € and mesh index (F).

### Pseudo-vascular networks exist over a range of time windows with varying levels of stability

In line with the qualitative estimation, the objects identified by quantitative morphological analysis are highly dynamic and diverse in their values across cell lines. The vectorial objects demonstrate nicely the structural variety of the pseudo-vascular networks formed by different cell lines. For example, for the 4T1-T cell line, total network length (Figure 5A) decreases over time, the number of the branches (Figure 5B) and extremities (Figure 5C) are relatively stable while the number of meshes (Figure 5D), total mesh area (Figure 5E) also decrease over time; conversely the mesh index (Figure 5F) increases to a peak before decreasing.

One might consider the total network length to be a good indicator of network stability, as similar general trends can be found over time across all cell lines tested, with a broad decrease over time in all cell lines, whereas other parameters show more divergent behaviour. Cell lines like MOC1 or MOC2, however, form pseudo-vascular networks that become smaller over time, possibly due to aggregation of the cells, which gives a false impression of their stability using the network length parameter. To examine stability of the networks, we therefore defined a stability score as the sum of the spread scores of all objects given time (Figure 6).

**Figure 2.**
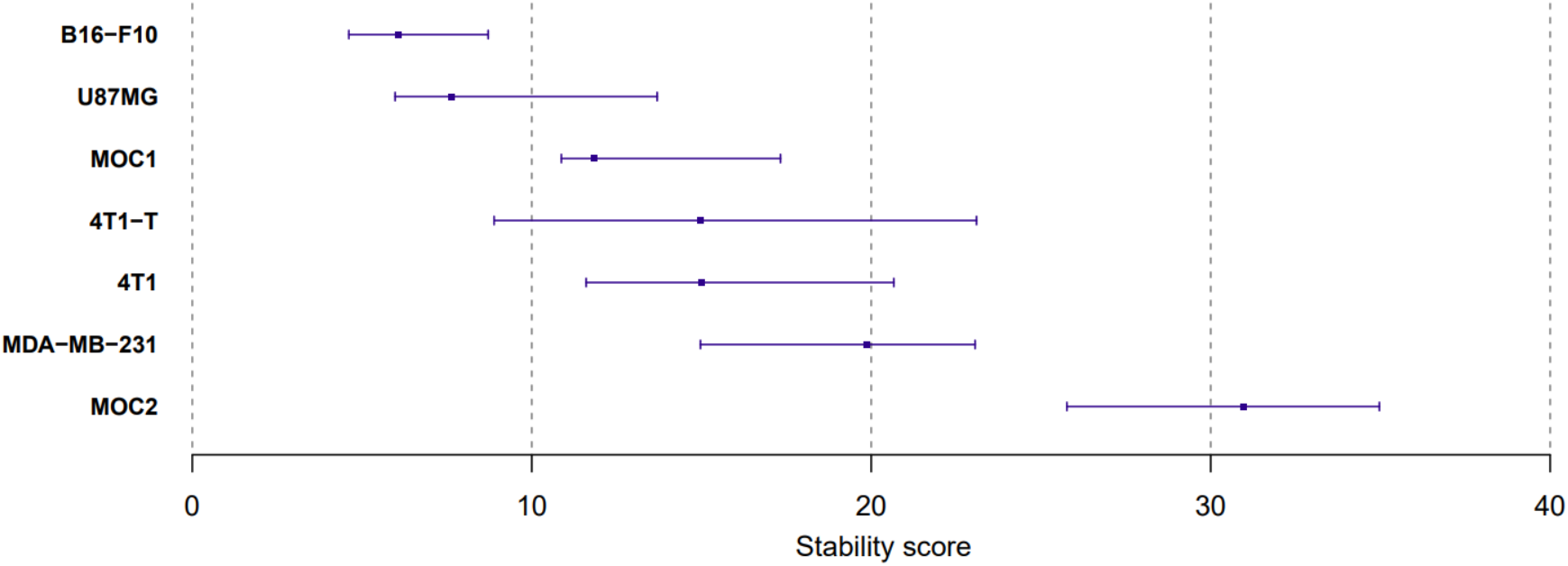
Cell line stability scores. Point and interval estimates of stability scores (x-axis) per cell line (y-axis) defined and 95% confidence intervals are shown. Inference was obtained by means of a hierarchical bootstrap. The level of uncertainty in stability scores (95% confidence interval width) is a consequence of the variability of the object values over time (see Supplementary Figure 1).

### Pseudo-vascular networks are differentially affected by the presence of drug treatment

We hypothesized that cell lines that exhibit a stable pseudo-vascular network phenotype would have stronger competence for VM. We next introduced perturbations to disrupt the networks. For the most stable of the cell lines studied, B16-F10, pseudo-vascular network evolution was examined under the presence of three FDA-approved agents previously reported to influence pseudo-vascular network formation [5]: multi-kinase inhibitor axitinib, the alkaloid brucine [26] and the mesenchymal-epithelial transcription factor inhibitor tivantinib [27] were tested against the DMSO solvent control.

We noted in initial testing that 0.1% DMSO affects the pseudo-vascular network evolution of all cell lines tested (Supplementary Figure 3), likely due to its effect on cell-to-cell and cell-to-matrix adhesion as DMSO can trigger cell aggregation [28]. Pseudo-vascular network inhibition has been reported before in endothelial cells for DMSO concentrations above 1% [29], however, the descriptive spatiotemporal analysis of this study shows that DMSO impacts cells even in lower concentrations and therefore, the tube assay protocol was optimized for 0.01% DMSO, that served as the control condition.

Pseudo-vascular network formation was impacted differentially by the presence of each of the drugs under study (Figure 7A). The effect of each drug on the mesh structures was then examined across a range of concentrations, as previously determined by the 2D drug screening assay, for the same timepoint (Figure 7B). Both the mean values of the vectorial objects and the variance were affected as compared to the control condition by all drugs tested. Tivantinib inhibits the formation of the meshes, even in dose an order of magnitude less than the two other agents, confirming prior studies that show this compound could be of interest in the context of VM.

**Figure 7.**
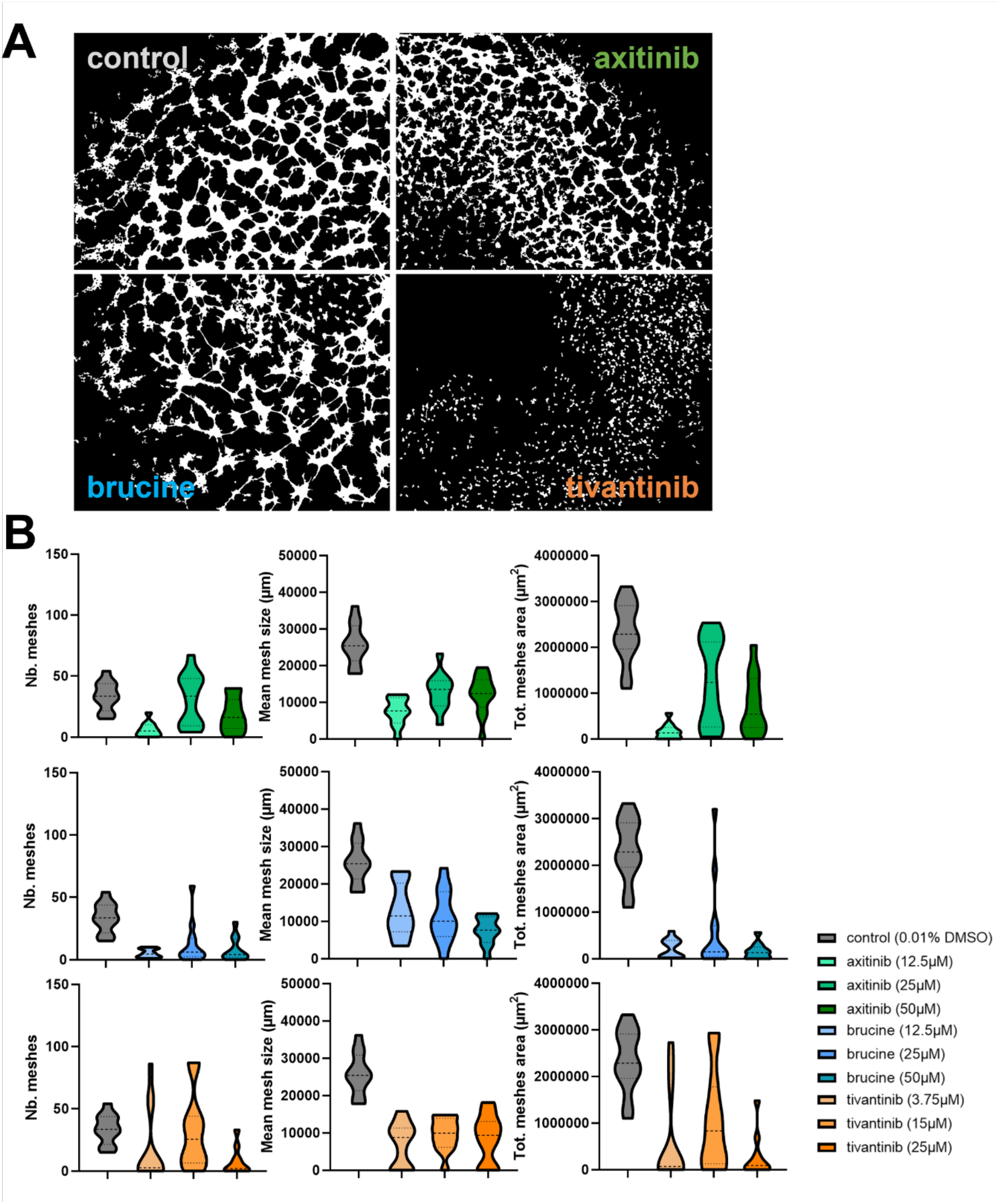
Drug screening assay demonstrating pseudo-vascular network disruption of different forms by compound. A. Binary masks of the B16-F10 cells forming tubular structures under the absence (control) or presence of 25μM of axitinib, brucine or tivantinib at the same timepoint (25hrs). B. The effect of each drug on the number of the meshes, the mean mesh size and the total mesh area within a range of concentrations.

## Discussion

To demonstrate the spatiotemporal dynamics of pseudo-vascular network formation and thus improve reproducibility of VM research, we report here a method of extracting pseudo-vascular network features *in vitro* that can be used to quantitatively compare findings from different studies. Representative cell lines from brain, breast, head and neck, and skin cancer types, that have been previously reported to express the VM phenotype [16, 30], were studied to compare their intrinsic VM competence. Furthermore, the sensitivity of these pseudo-vascular networks to different therapeutic compounds was studied to compare their effect on network formation and stability.

It is well known that endothelial cells are conditionally able to form a structured physiologic-like network that remains stable over time *in vitro*, but to understand the spectrum of VM capabilities across cancer cell lines, our findings demonstrate that it is important to follow their network dynamics rather than making measurements at a single time point. Repurposing Angiogenesis Analyser, an open-source toolkit that is designed to describe endothelial morphologies, we extracted pseudo-vascular vectorial object descriptors to create a statistical framework to compare VM in cancer cell lines over time, normalized to account for proliferation differences. The comparative study performed led to a complete framework to: i) estimate the VM competence of different cell lines; ii) describe the *in vitro* morphology of the pseudo-vascular VM networks, irrespective of spatiotemporal differences; and iii) enable the identification of potential anti-VM chemotherapeutic agents.

Between the seven cell lines used for the purposes of this study, the pseudo-vascular networks formed were spatiotemporally distinct. For all cell types, objects related to the number of the meshes formed and the total mesh area, but not the mean mesh size, were able to discriminate the different pseudo-vascular networks over time. Meshes are a combination of the nodes and the connected branches in the pseudo-vascular networks, a formation that has been reported in both the tubular and the patterned matrix type VM [2, 5]. B16-F10 and U87MG cell lines depict the lowest variance among all the network descriptors examined here, unlike the MDA-MB-231, which persist for a longer time window. Although B16-F10 and 4T1-T are highly proliferative cell lines, the stability of their VM pseudo-vascular networks is high, which suggests that VM competence is independent of the proliferative capacity of the cells. Additionally, our methodology was successful in showing subtle differences between 4T1 and 4T1-T derivative cell line related to their VM competence [7].

Having established relevant vectorial objects and measures to discriminate between cell lines, the most stable line (B16-F10) was then subjected to a screening of three anti-vascular agents that are mechanistically-related to VM [5]. Although none of the drugs tested appeared to be cytostatic or cytotoxic for the B16-F10 cell line, as no change in the confluence was observed between treatment and control conditions, all of them were able to lead to irreversible pseudo-vascular network disruption in very low concentration. Furthermore, vectorial objects related to the mesh structures were sufficient to estimate the impact of the inhibition of the drugs to the pseudo-vascular network architecture. Our method could be applied in future to screen chemotherapeutic regimens exhibiting a dual anti-vascular and cytotoxic effect in VM-competent cancer cells [31, 32].

We have demonstrated potential for dynamic imaging to inform on the diversity of VM-competence between cancer cells that form pseudo-vascular networks *in vitro*, however, there remain several limitations to our approach that should be addressed in future. First, cancer cells form tubes in the xy plane of the BME-coated wells used here and fail to form 3D structures [13, 33]. VM tubular structures are often formed by one layer of cancer cells that may or may not be a functional tubule and this process is cell line-specific [34]. The application of more advanced cell culture protocols with confocal microscopy may enable 3D tubular structures to be studied using a similar approach. In addition, the BME-coated wells lead to a meniscus effect, which required some data pre-processing and excluded some of the potential vectorial objects from being applied due to imaging artefacts. Improved preparation to minimise the meniscus could be developed.

A second limitation is that we examined the intrinsic capacity of the cancer cells to form tubular structures in the absence of any supportive cells of the tumour stroma or the host. Thus, the homology to real vascular networks is low and cannot substitute the complexity of the *in vivo* tumour vasculature, focusing instead on the elementary VM structures. Nonetheless, the same experimental framework, adjusted for optimal cell seeding density and monitoring time, could also be used in co-culture settings and/or pre-conditioned medium in order to further study the molecular basis of the VM biological phenomenon [25].

A final limitation concerns the Angiogenesis Analyser Fiji plugin. The plugin has been used for the analysis of networks other than those formed by HUVEC endothelial cells, such as the mouse retina vascular network captured in fluorescent microscopy images [35], depicting its wide applicability in other forms of networks and imaging data. However, the skeletonization process that is applied by the Angiogenesis Analyzer is not able to characterize wider tubular formations as those expected in a patterned VM type [9], which may require further algorithm developments to enable the full range of VM phenotypes to be characterized.

## Conclusions

In this study, we presented a framework for the *in vitro* assessment of VM that includes a generic tube formation assay that allows the longitudinal monitoring of the pseudo-vascular VM networks and the application of an open-source software plugin for the quantitative morphometric description of the networks. We apply this workflow to a relative comparison of reported VM-competent cell lines and evaluate the effect of three clinically-applicable anti-vascular drugs on network formation. Our findings indicate that the meshed structures of the pseudo-vascular networks can serve as a potential VM *in vitro* marker irrespective of the cancer type. The proposed methodology can be further applied accounting for more cell lines and/or the anti-VM efficacy of therapeutic schemes.

## Supporting information

Supplementary video 2 MDA-MB

Supplementary video 5 MOC1

Supplementary video 6 MOC2

## Acknowledgements

The authors would like to thank CRUK-CI Core Facilities for their support and especially Manuela Natoli from RICS. This work was supported by Cancer Research UK (C9545/A29580, C47594/A29448, C17918/A28870). For the purpose of open access, the authors have applied a Creative Commons Attribution (CC-BY) license to any Author Accepted Manuscript version arising.

## Author Contributions

MEO designed, performed and analyzed the biological experiments, as well as guided the statistical analysis and wrote the paper manuscript. DLC designed and implemented the statistical analysis, as well as contributed as a writer to the paper manuscript. EVB co-performed and analyzed part of the biological experiments, as well as co-designed the drug screening assay and contributed as a writer to the paper manuscript. IC provided part of the *in vitro* protocols, as well as the 4T1 and 4T1-T cancer cell lines. MG co-performed part of the biological experiments. GH supervised the biological experiments. VS co-designed and supervised the biological analysis. SEB co-conceived and supervised the project, evaluated the results and contributed as a writer to the paper manuscript. All authors have read and approved the final manuscript.

## Data availability statement

All data and analysis scripts associated with the manuscript will be made available upon publication of the paper at: https://doi.org/10.17863/CAM.101798

