## Supplementary information for "The *in vitro* dynamics of pseudo-vascular network formation"

vasculogenic mimicry, tube formation assay, pseudo-vascular networks, morphometric analysis, descriptive statistical model

**Supplementary Figure S1**

Selected vectorial objects of the pseudo-vascular networks of the different cell lines over time, as described by the linear regression estimator.

## 1a - U87MG

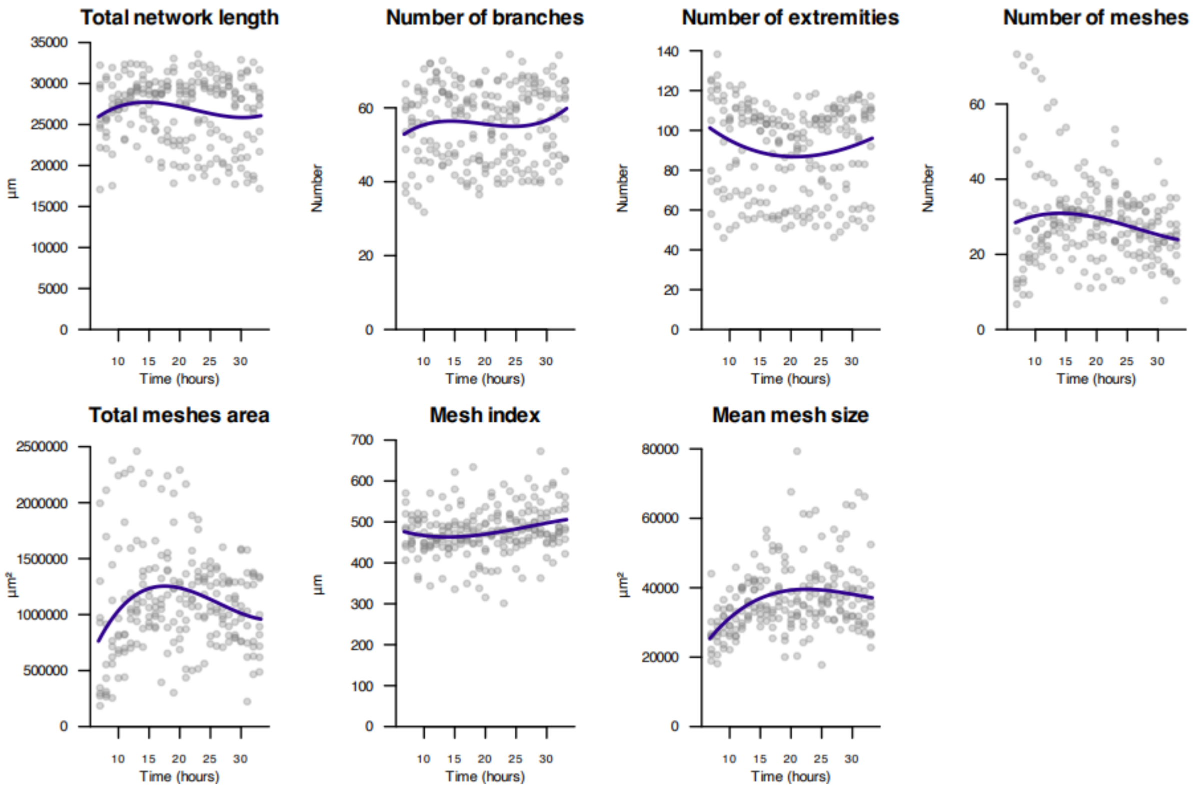

### 1b - MDA-MB-231

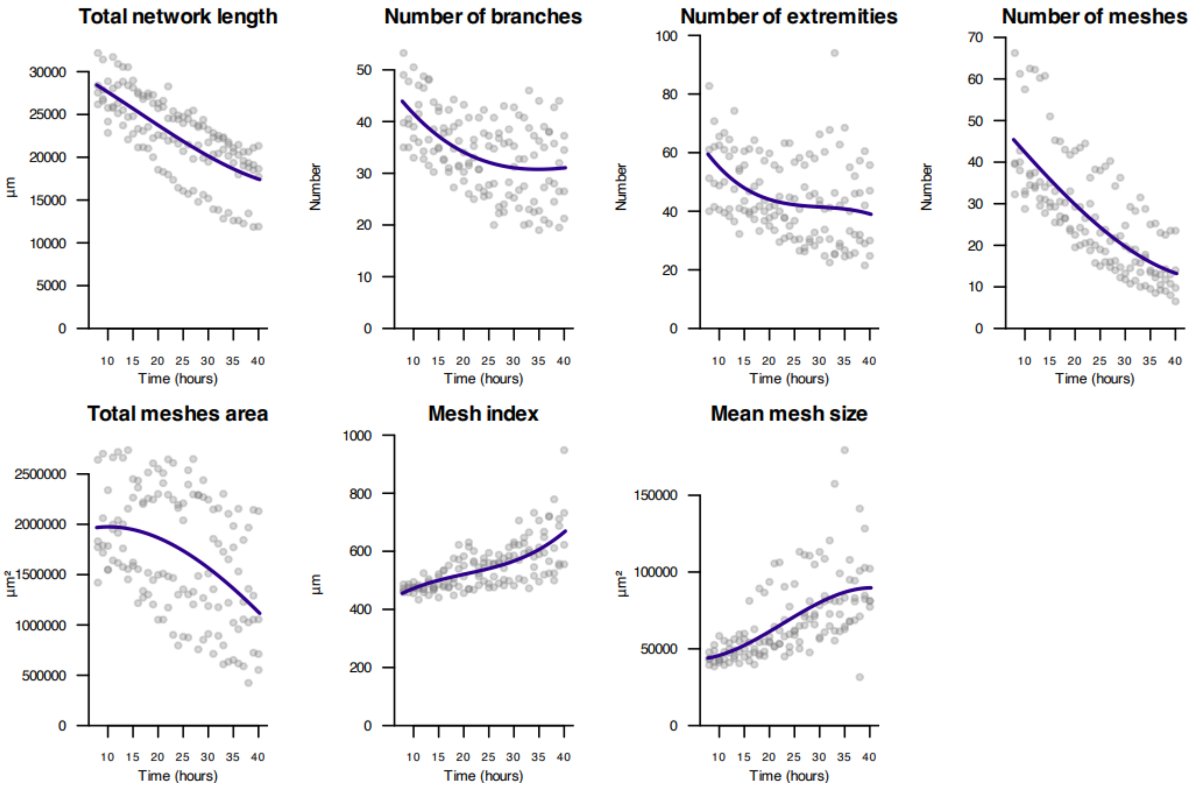


## 1c - 4T1

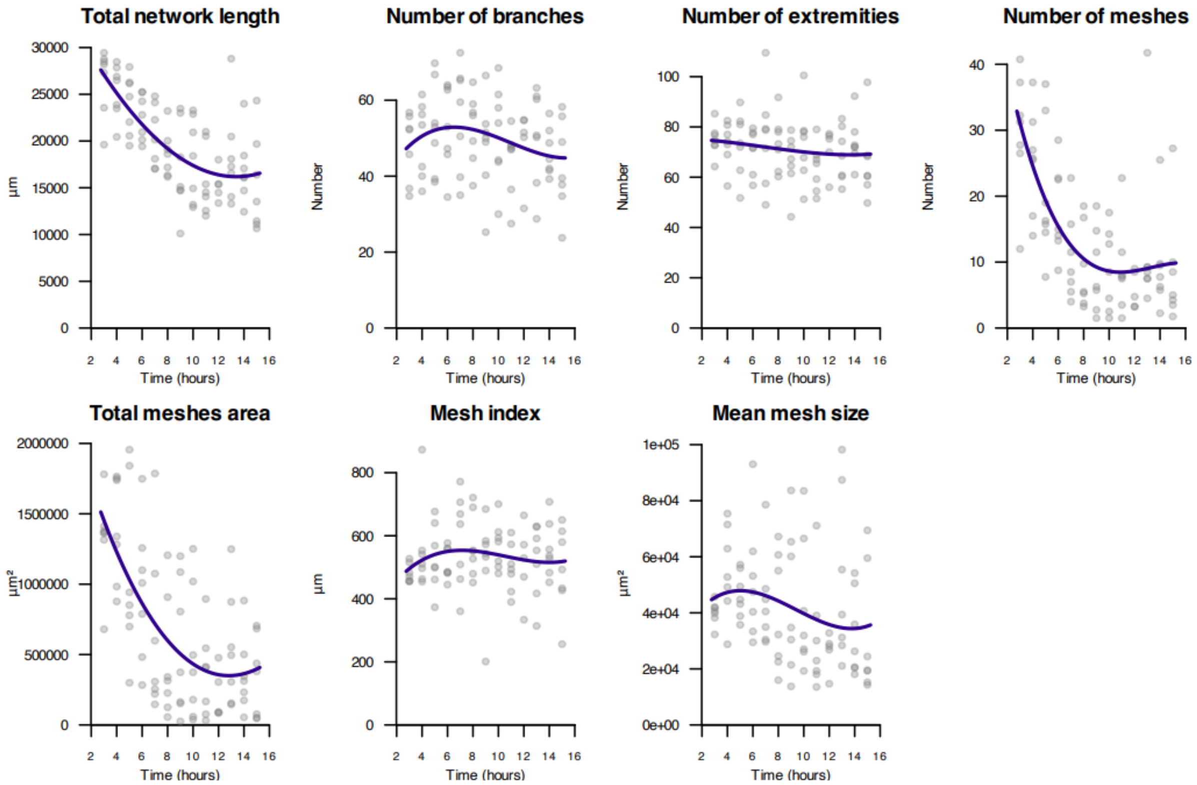

# 1d - 4T1-T

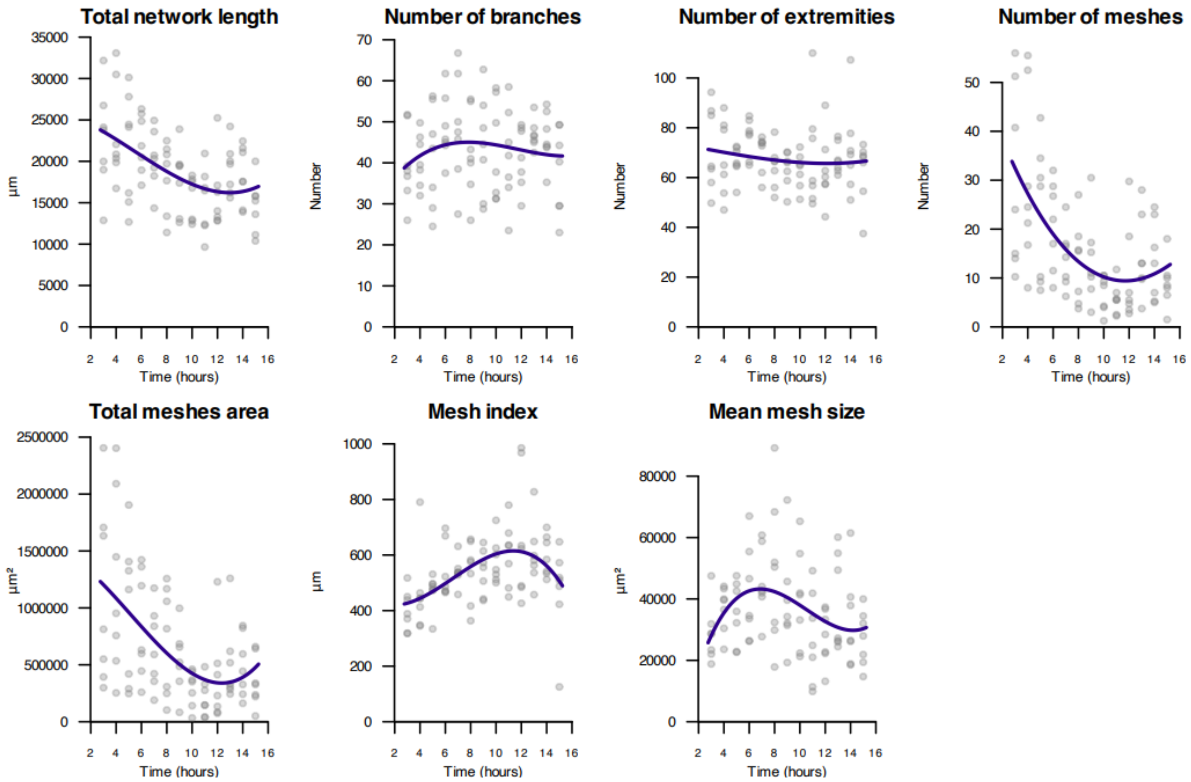


# 1e - B16-F10

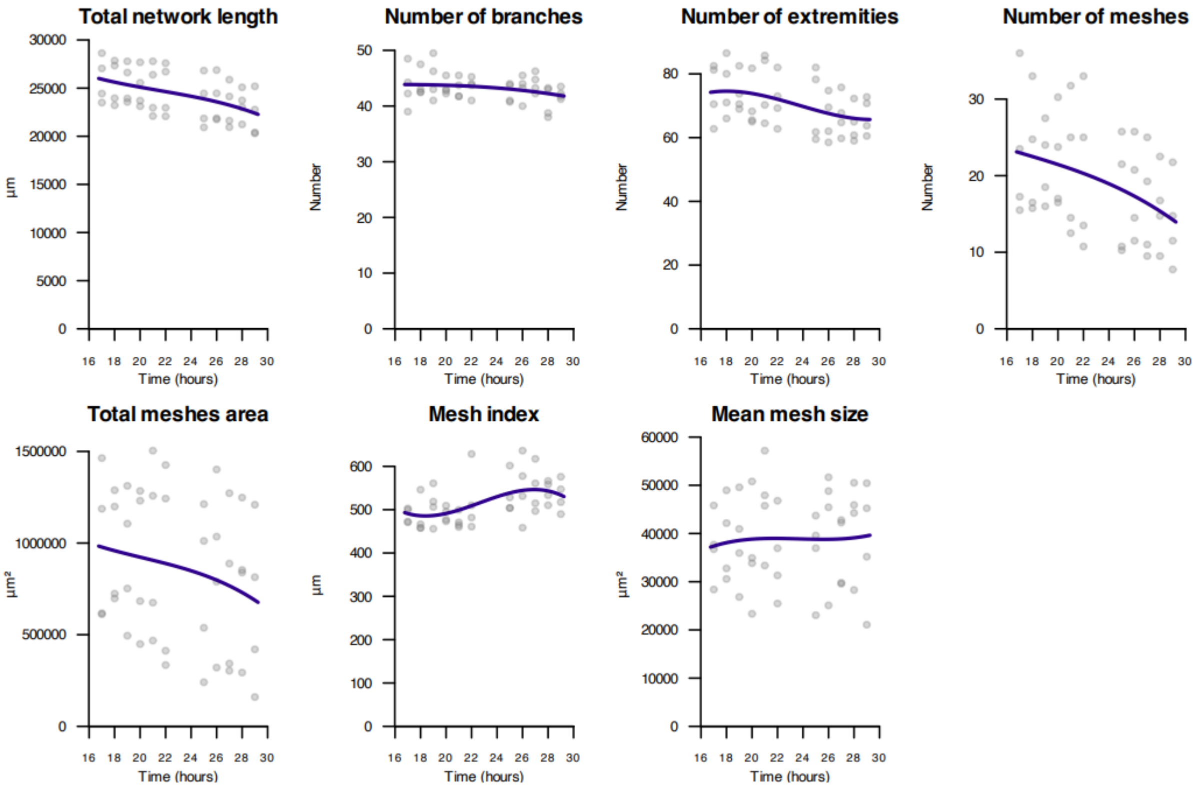

### 1f - MOC1

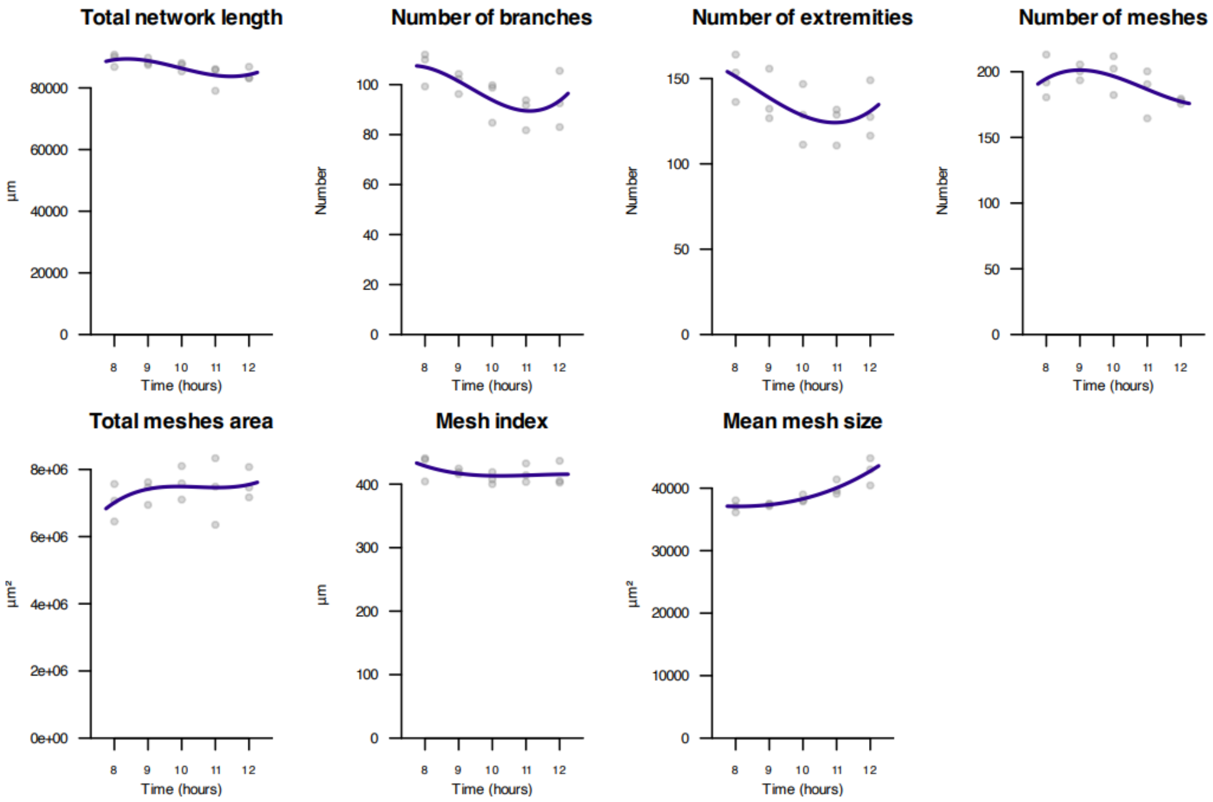


### 1g - MOC2

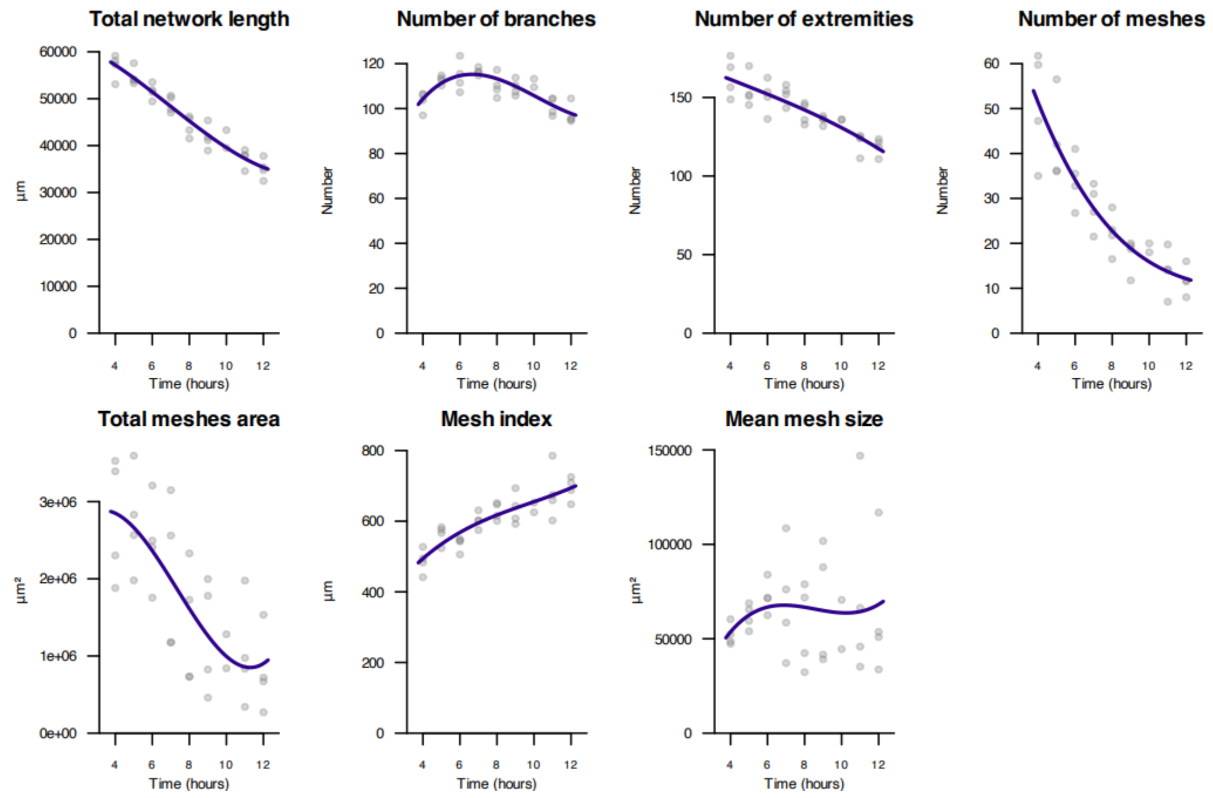

### Supplementary Figure S2

Stability scores for each of the vectorial objects as described by the linear regression estimator applied to each individual feature and cell line (95% confidence interval width).

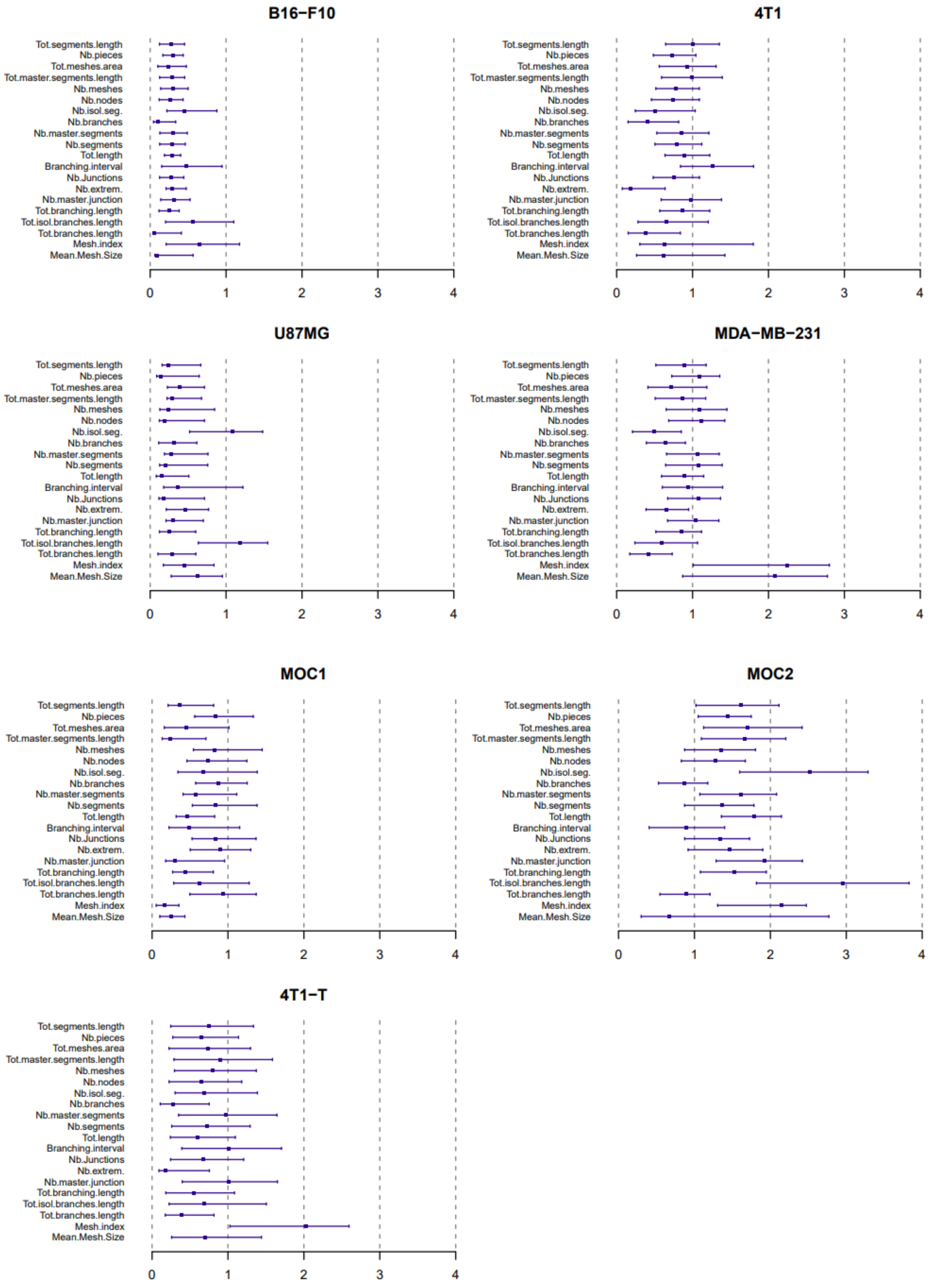

**Supplementary Figure S3**

The effect of the DMSO on the meshes of the pseudo-vascular networks of 4T1 and 4T1-T cell lines. DMSO concentrations other than 0.1% have been tested because of the effect of the DMSO on the tubular capacity. The 0.01% DMSO concentration seems to recapitulate the absence of any vehicle (control) more closely and therefore, it was used for the drug-treated tube formation assay.

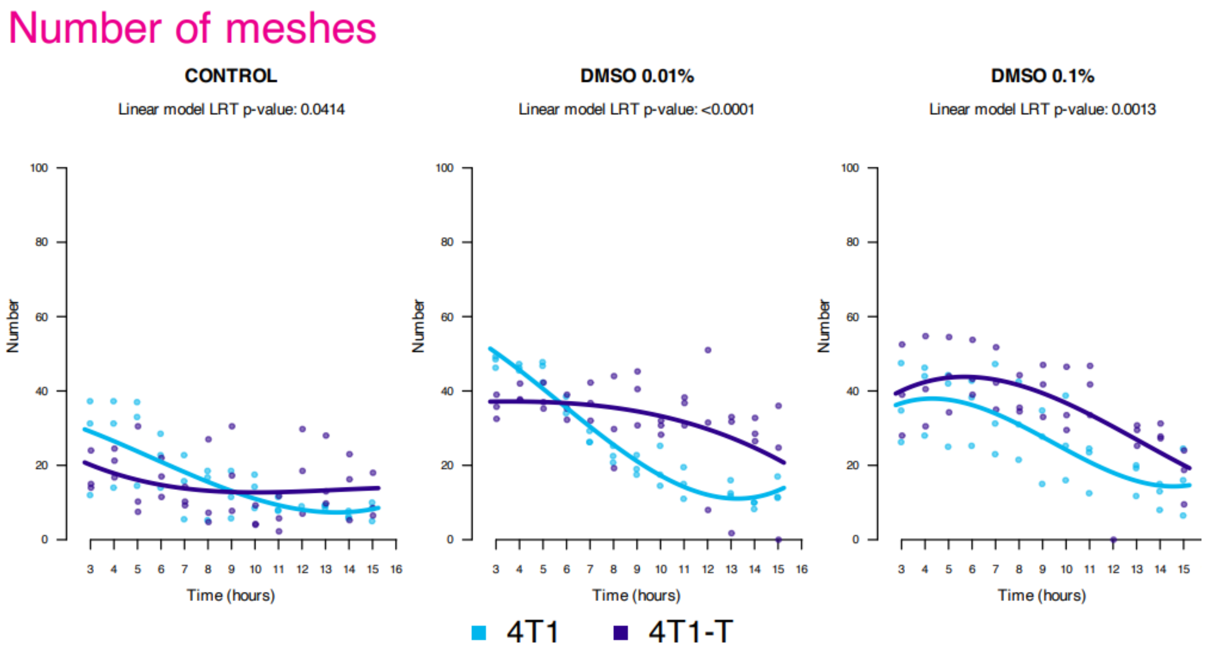

**Supplementary Table 1**

Definition of the pseudo-vascular network vectorial objects as defined in [19].

| Vectorial Object | Definition | Morphological Feature |
|---|---|---|
| **extremity** | ***pixel with only one neighbour*** | number of extremities |
| **node** | ***pixel with at least 3 neighbours*** | number of nodes |
| **junction** | ***group of dots forming a bifurcation*** | number of junctions<br>number of master junctions |
| **segment** | ***binary line linked with two junctions***<br><br>***(a piece is a segment intercepting a circle)*** | number of segments<br>number of master segments<br>total master segments length<br>number of isolated segments<br>total segments length<br>number of pieces |
| **branch** | ***line linked to one junction and one extremity*** | number of branches<br>total branching length<br>total branches length<br>total isolated branches length<br>branching interval |
| **mesh** | ***closed area formed by segments and their junctions***<br><br>***index: total master segments length / number of master segments*** | number of meshes<br>total mesh area<br>mean mesh size<br>mesh index |
| **network** | ***sum of length of segments, isolated elements and branches*** | total length |

**Supplementary Videos**

A sequence of phase-contrast images in 4x magnification, focusing on one quarter of a representative well for each of the cell lines, in order to monitor the cells undergoing the tube assay. Time slightly varies depending on the example, up to ~3 days post seeding. Acquisition is every 1hr and scale bar is set to 1mm. Videos have been made using the Incucyte software. Plate lid condensation, lost timepoints and changes in the focal plane can be expected during the automated process, as the experiments run uninterrupted for the whole time course.

Pseudo-vascular network time window is within this time period, but is always concluded before 2 days for most of the cell lines and is often followed by a cell aggregation period. Changes in the confluence of the cells can be observed in the later phase of the time window, indicating proliferation of the cells after they are organized in tubes. Especially for the MOC2 cell line, there is evidence of proliferation (differentiation of the cellularity and the diameter of the tubes) during the organization into the pseudo-vascular network, but this is beyond the scope of this study.

**Video S1 –** U87MG

**Video S2 –** MDA-MB-231

**Video S3 –** 4T1

**Video S4 –** 4T1-T

**Video S5 –** MOC1

**Video S6 –** MOC2

**Video S7 –** B16-F10
